# Beak morphometry and morphogenesis across avian radiations

**DOI:** 10.1101/2023.02.21.529429

**Authors:** Salem Mosleh, Gary P. T. Choi, Grace M. Musser, Helen F. James, Arhat Abzhanov, L. Mahadevan

## Abstract

Adaptive avian radiations associated with the diversification of bird beaks into a multitude of forms enabling different functions are exemplified by Darwin’s finches and Hawaiian honeycreepers. To elucidate the nature of these radiations, we quantified beak shape and skull shape using a variety of geometric measures that allowed us to collapse the variability of beak shape into a minimal set of geometric parameters. Furthermore, we find that just two measures of beak shape, the ratio of the width to length and the normalized sharpening rate (increase in the transverse beak curvature near the tip relative to that at the base of the beak), are both strongly correlated with diet, and thus indicative of how bite forces are correlated with beak shape. Finally, by considering how transverse sections to the beak centerline evolve with distance from the tip, we show that a simple geometry-driven growth law termed “modified mean curvature flow” captures the beak shapes of Darwin’s finches and Hawaiian honeycreepers. A surprising consequence of the simple growth law is that beak shapes that are not allowed based on the developmental program of the beak are also not observed in nature, suggesting a link between evolutionary morphology and development in terms of growth-driven developmental constraints.

## 1 Introduction

Avian adaptive radiations, such as Darwin’s finches and Hawaiian honeycreepers, provide beautiful examples of how one species can diversify into many ecologically distinct ones, with the beak evolving into a variety of forms that enabled new functions. To elucidate this diversification of form, we need to quantify both the extent and nature of variation in beak shapes and how this variation enables function (e.g., feeding and vocalization). In addition, this variation in shape is generated by modifications of a developmental program to produce new forms, and it is important to describe these three aspects of beak shape evolution (form, function, and development) in a common framework that can further elucidate the nature of the adaptive radiation. Here, using micro-computed tomography (*μ*CT) scans of Darwin’s finches, Hawaiian honeycreepers, and their relatives, we quantify beak and skull shape variation to elucidate their evolution and development.

Beak shapes have been quantified using discrete measurements, such as length, width, and depth, or using a set of landmarks — identifiable points that are common to all studied specimens [1, 2]. While these methods are valuable in elucidating beak shape variation [3–5], they do not explicitly capture the rich information in beak shape, such as the curvature of its surface. Here, to fully describe beak shape, we quantified its upper surface using three complementary approaches. In the first approach, we extract the upper bill centerline and measure its length and curvature, in addition to extracting cross sections of the beak normal to the centerline. The second approach uses a functional form that accurately describes the upper surface of the beak, where the beak shape is represented by transverse parabolas with curvature linearly decreasing with distance from the tip [6]. The third approach employs the Hausdorff distance [7] to quantify how each pair of beaks (or pair of skulls) are different from each other. After using these methods to construct morphospaces for beak and skull shapes, we explore patterns of beak shape variation and their relation to diet and mechanical function.

Motivated by studies of beak growth during embryonic development [8, 9], where a group of proliferating cells near the tip of the developing beak form a growth zone that defines the shape of the beak as it shrink over time, we investigate how transverse beak cross sections normal to the centerline change as they approach the tip of the beak. We observed that beak cross sections change shape over time, becoming more convex and circle-like the closer they are to the tip, which is contrary to the previous assumption that the growth zone will shrink uniformly (without change in shape) over time [9]. This observation motivates us to study a different model of growth zone evolution, where a highly curved region near its boundary shrinks (stop dividing) at a higher rate compared with cells in the bulk of the growth zone. This tissue scale behavior can emerge from a morphogen diffusion and degradation, where cells continue dividing if the concentration of this morphogen is above a threshold value [6]. By additionally allowing changes in the growth direction that can generate beaks with highly curved centerlines, we are able to generate the observed beak shapes of both Darwin’s finches and Hawaiian honeycreepers.

## 2 Evolutionary morphospace of beaks

### 2.1 Constructing morphospaces

In Fig. 1, we show beak meshes from Honeycreepers (n = 41) and their relatives (n = 9) along with their phylogenetic relationships (see SI Fig. S1 for views of the entire skull). Our dataset, with 151 total specimens, also includes Darwin’s Finches (n = 54) and their relatives (n = 47). To quantify their 3-dimensional shapes, we extract and smooth the 3D surfaces of the upper beaks from *μ*CT-scans of the skulls and, for each smoothed beak mesh, automatically extract its centerline and beak cross sections normal to it (see Fig. 1B and SI Figs. S2 for details of smoothing and centerline extraction).

**Figure 1:**
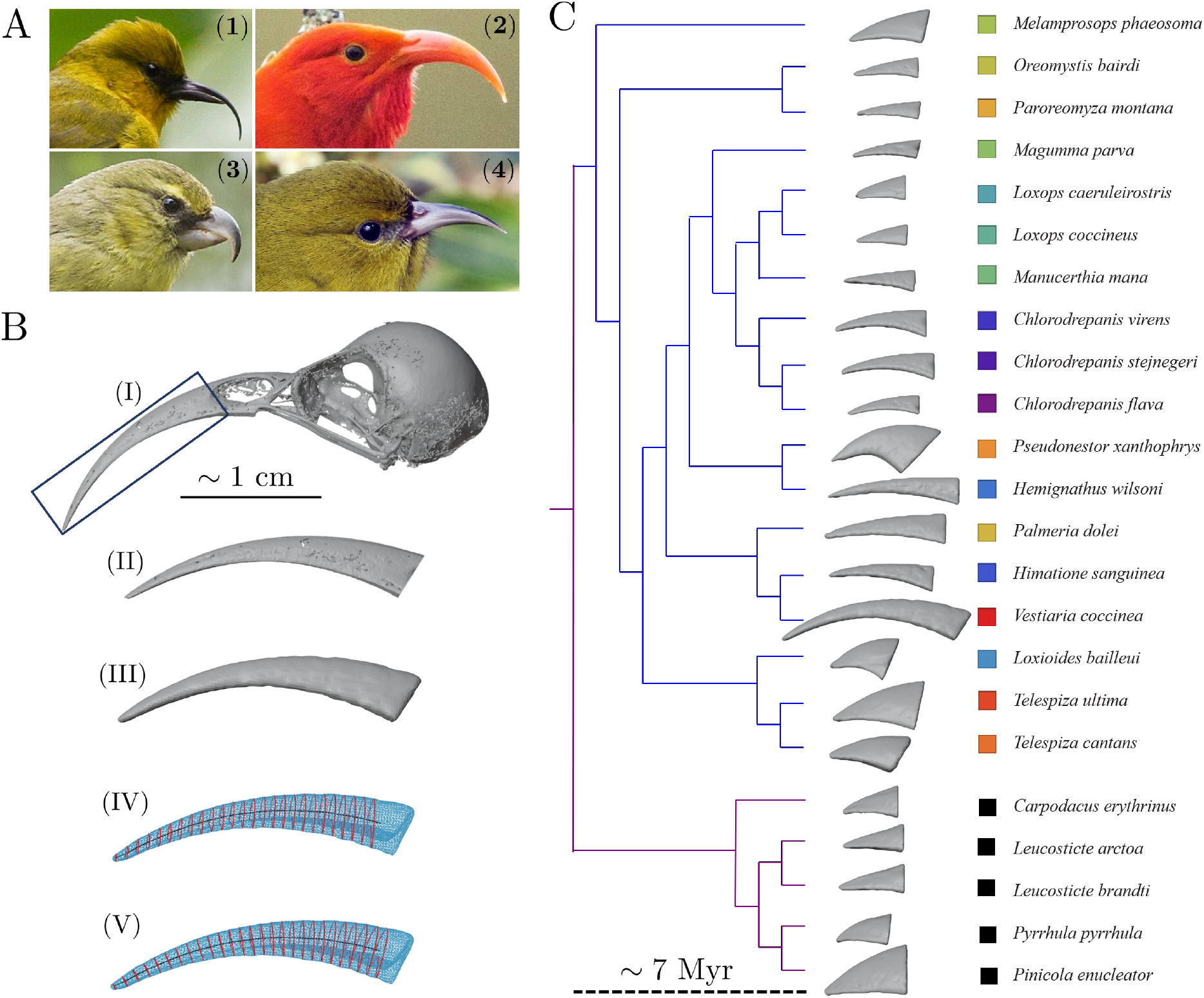
Beak morphology and phylogeny of Hawaiian honeycreepers. (A) Examples showing the diversity of beak shapes observed for Hawaiian honeycreepers: (1) *H. wilsoni* (2) *V. coccinea*, (3) *P. xanthophrys*, and (4) *C. flava*. (B) Steps used in our analysis to extract beak shapes, centerlines and cross sections (see SI for details). (I) The skull is aligned so that its long axis is in the x-direction. (II) The beak is cut from the skull and (III) smoothed to remove holes from the mesh. (IV) starting with vertical cross sections perpendicular to the x-axis, we find a test centerline as the center of mass (assuming uniform density) for each cross section. New cross sections are obtained normal to the generated centerline. Iterating this procedure we get the final centerline and cross sections shown in (V). (C) The phylogeny of honeycreepers and their relatives based on Ref. [10].

To extract the upper surface of the beak we first find the tomium (cutting edge of the beak) as the points with minimum and maximum lateral coordinates (y-axis in Fig. 2A). The tomium then separates the upper and lower surfaces of the beak, which we orient so that the origin of the coordinate system is at the tip of the upper surface, with axes oriented to correspond to the principal axes (SI Fig. S3). We find that the following paraboloidal profile, which was introduced as a fit to the beaks of Darwin’s finches in [6], captures the upper surface of the beak for the samples in our dataset (SI Fig. S3):

**Figure 2:**
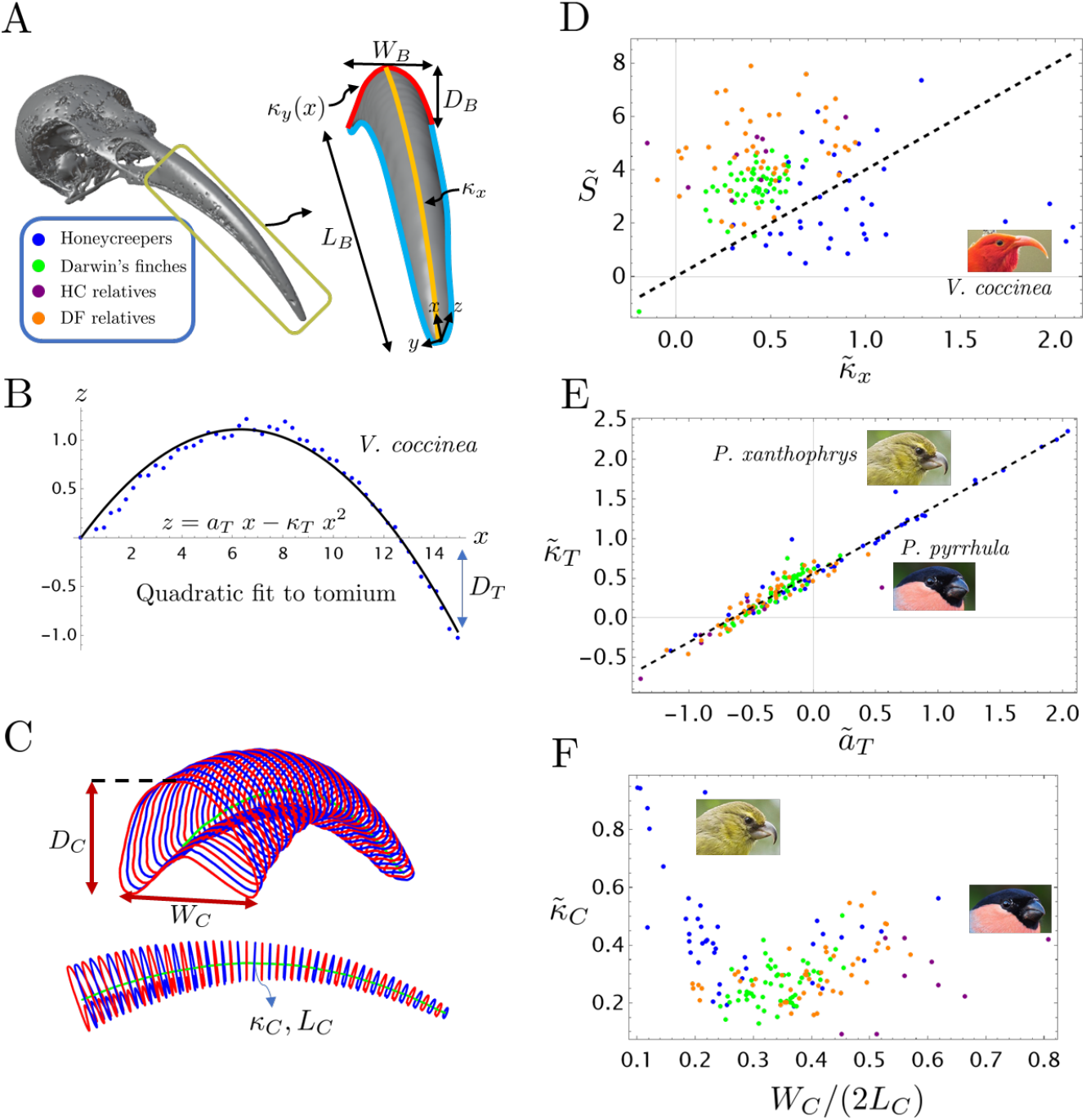
Evolutionary morphospace of beaks. (A) The extracted upper beak (right) from a V. *coccinea* skull (left). The beak axes are aligned with its principal directions (Fig. 1B). The functional form 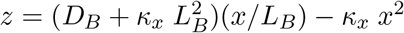 approximates the midsagittal curve (*y* = 0) shown in yellow. Transverse sections (red) also have parabolic form with curvatures varying with distance from the tip according to *κ_y_*(*x*) = *κ_tip_* − *S x*. The tip curvature, *κ_tip_*, is extrapolated from the relation *κ_tip_* ≡ *κ_y_*(0). (B) The extracted tomium of the beak projected onto the *xz*−plane and the corresponding parabolic fit. (C) The morphological variables defined by the beak centerline, where *L_C_* is its length and *κ_C_* is its curvature. At the base of the beak, the cross section is characterized by its depth *D_C_* and width *W_C_*. (D) Beak morphospace defined by the variables 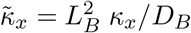 and 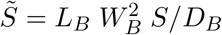 for 151 specimens from 4 groups, color coded as shown in panel (A). (E) The morphospace defined by the dimensionless tomium parameters 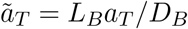 and 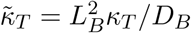. The fact that most points are close to the dashed curve, indicates that *D_T_* ≈ 0.5*D_B_* for almost all beaks in our dataset. (F) Beak morphospace defined by the aspect ratio *W_C_*/(2*L_C_*) and dimensionless centerline curvature 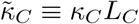.

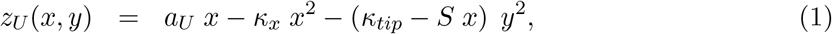

where the subscript *U* denotes the upper surface, *a_U_* gives the slope of the midsagittal section (*y* = 0) near the tip, *κ_x_* is the curvature of the midsagittal section, *κ_tip_* is the curvature in the transverse direction at the tip and *S* represents the “sharpening rate” of the beak curvature towards the tip. Since the size of the cross section shrinks to zero at the tip, the parameter *κ_tip_* is extracted from the linear fit of the transverse curvature *κ_y_* ≡ *κ_tip_* − *S x*. The beak is also characterized by its length *L_B_*, width *W_B_*, and depth *D_B_*, where the subscript B denotes measurements aligned with the bounding box of the beak oriented along the principal directions (Fig. 2A). To compare beak shape across species, we remove the effect of scale and consider the dimensionless shape variables 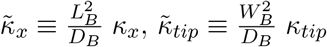 and 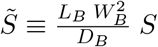.

To describe the shape of the tomium, we project the extracted tomium points onto the *xz*-axis and fit the resulting curve to a parabola,

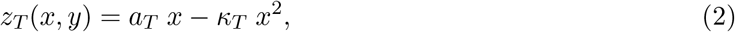

which gives a good fit as shown in Fig. 2B. The 3D tomium curve can then be found as the intersection of the upper beak surface given by Eq. (1) and the surface given by Eq. (2). We also define normalized tomium parameters as 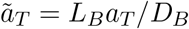 and 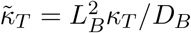. Knowing the upper beak surface and tomium, Eqs. (1)-(2), we can define the beak depth and width as a function of the coordinate *x*. The depth is defined as *D*(*x*) ≡ *z_U_* (*x*, 0) − *z_T_* (*x*, 0), while the depth is defined through the equation *z_U_* (*x, W* (*x*)/2) = *z_T_* (*x, W* (*x*)/2), which leads to

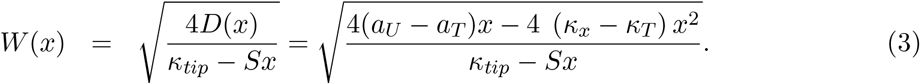

Comparisons of this equation with data is given in SI Fig. S3.

To elucidate the role of centerline curvature in honeycreeper beak shape variation, we compute the arc-curvature of the beak centerline, *κ_C_*, by fitting the centerline to a circular arc (Fig. 2C and SI Fig. S3). In addition, as shown in Fig. 2C, we define centerline adapted length, width and depth (*L_C_, W_C_, D_C_*) and corresponding centerline adapted coordinates (*x_C_, y_C_, z_C_*), where *x_C_* is the arc-length coordinate along the beak centerline and (*y_C_, z_C_*) represent the remaining two orthogonal directions. We also define a dimensionless curvature 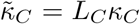 and aspect ratio *W_C_/L_C_* between the width of the beak basal cross section and the centerline length.

Lastly, to elucidate the statistical distribution of beaks in morphospace, and the covariation between beak and skull shapes, we perform a surface-based morphometric analysis for the beaks and skulls in our data set. We first normalized each surface mesh based on the distance between the beak tip and the centroid of the mesh. Then, for each pair of specimens, we searched for an optimal rigid transformation (i.e. a combination of translations and rotations) to align them and compute the symmetric Hausdorff distance between them [7, 11]. Once we obtained the distance measure for all pairs of specimens, we use multidimensional scaling (MDS) to represent all specimens on the 2D plane, where distance between points in this plane are as close as possible to the computed Hausdorff distance between each pair of meshes [12, 13]. Repeating this for beaks and skulls separately, results in coordinates that describe the beak and skull shapes. In addition, by using affine (rigid, scaling and shear) transformations to optimally align meshes, we obtain MDS coordinates that indicate how well two shapes can be transformed into each other with this set of transformations [14].

### 2.2 Patterns in morphospace

By looking at the samples in the morphospace 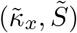 (Fig. 2D), we note that the region 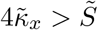 is only occupied by Honeycreepers. The constraint 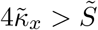, which is satisfied by all the other species, was predicted in [6] as a developmental constraint resulting from curvature driven growth of the beak. The fact that Hawaiian Honeycreepers do not satisfy this constraint is related to their higher normalized centerline curvature 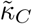 (Fig. 2F) and large midsagittal curvature 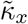 coupled with relatively low sharpening rates (Fig. 2D).

Since the curvature of the parabola at the base of the beak is given as 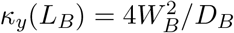, we have the identity 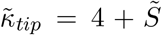, which we verify by calculating the Pearson correlation coefficient between the two quantities across our 151 samples 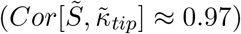 and noting that specimens are close to the plane 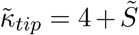 in morphospace (SI Fig. S4A). This reduces the morphospace of beak shapes to an overall size (which may be taken as the length *L_B_*), two aspect ratios associated with the relative depth *D_B_/L_B_* and width *W_B_/L_B_* and two scaled curvature parameters 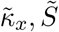.

Looking at the normalized tomium parameters (Fig. 2E), we find that points lie close to the line 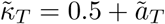. To understand what this means we note that the depth of the tomium below the tip, denoted by *D_T_* in Fig. 2B, can be calculated using Eq. (2) as 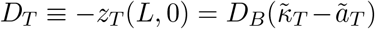. Therefore, the line in Fig. 2 E implies that *D_T_ /D_B_* ≈ 0.5 for most specimens in our dataset, with the Maui parrotbill (*P. xanthophrys*) and Eurasian bullfinch (*P. pyrrhula*) deviating from this trend due to their extreme morphology (see also Fig. 2F).

We observed that the depth over width aspect ratio *D_C_/W_C_* for the most basal cross section normal to the centerline is nearly constant for Honeycreepers (*D_C_/W_C_* = 0.64 ± 0.1, where 0.64 is the mean and 0.1 is the standard deviation), which is close to the value for Darwin’s Finches (*D_C_/W_C_* = 0.68 ± 0.07), as can be seen in SI Fig. S4B. In addition, the dimensionless curvature 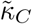 is large for Honeycreepers, especially for *V. coccinea*, as expected (Fig. 2F). However, this high value of dimensionless curvature is driven more by its longer length *L_C_* than the absolute value of its curvature *κ_C_* relative to other species (SI Fig. S4C).

As can be seen from Figs. 2D-E and 3A (see also SI Fig. S4), Honeycreepers, Darwin’s finches and their respective relatives occupy distinct regions of morphospace, with Honeycreepers occupying a broader and unique range of variation in each case. This observation is also confirmed using the beak MDS coordinates from the surface based analysis using rigid transformations (Fig. 3A). By contrast, the overlap between the regions occupied by the different groups in the full skull MDS plane is more significant (Fig. 3B). These results are consistent with the those presented in [5], in which the principal component analysis (PCA) was applied to a set of normalized landmarks representing the beaks or the full skulls. To explore further the nature of variation between the different groups, we consider MDS coordinates based on affine matching for both beaks and skulls. For beaks, we find that Darwin’s finches occupy a distinct region, indicating that as a group, we cannot match their beaks with affine transformation to the other groups in our study (Fig. 3C).

**Figure 3:**
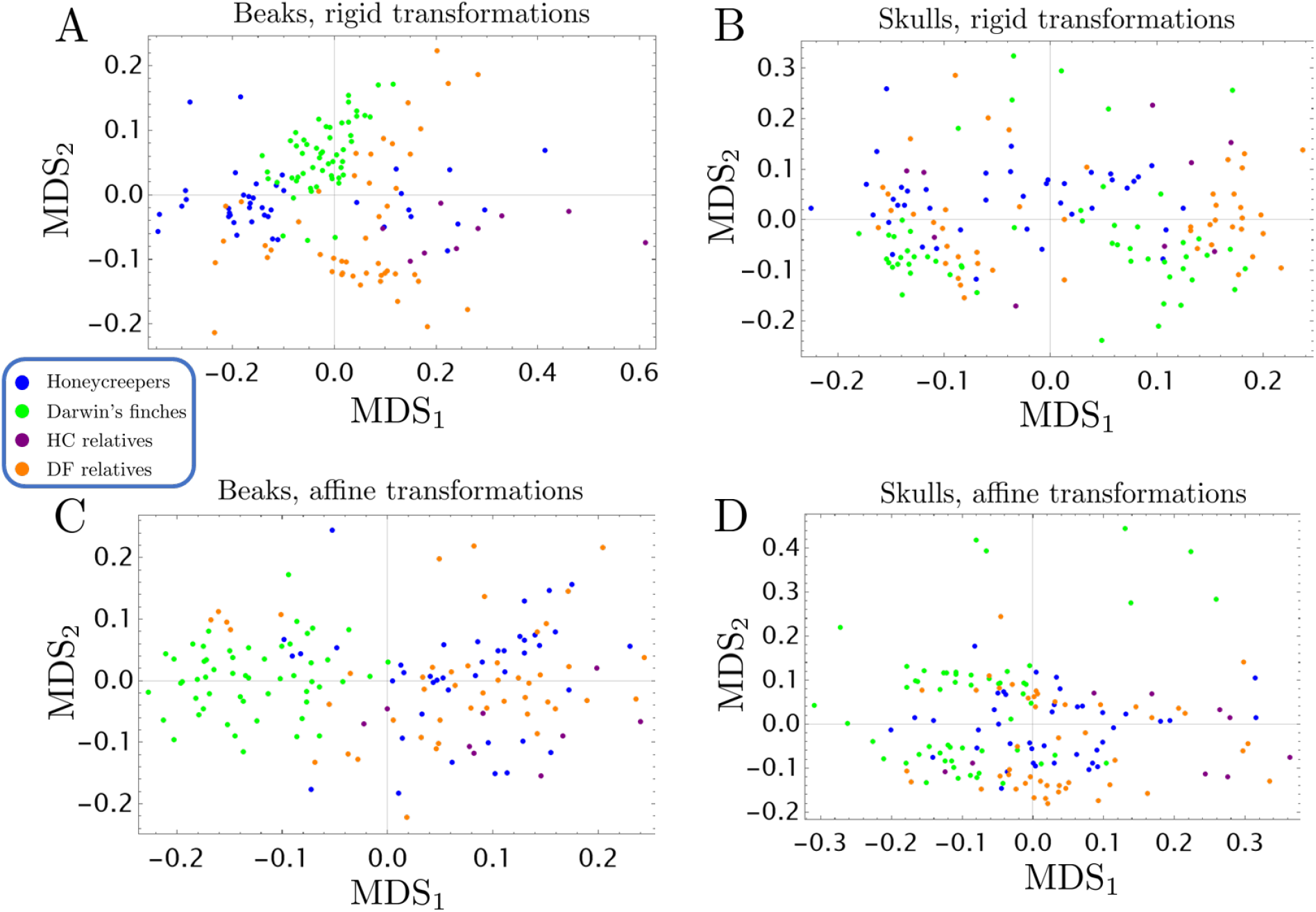
Multidimensional scaling (MDS) results of the surface-based morphometric analysis for the beaks and skulls. (A) Coordinates calculated using MDS analysis of the symmetric Hausdorff distance between beak meshes after optimally aligning them (with respect to the Hausdorff distance) using a rigid transformation. (B) Same as (A) but for the entire skull. (C) Same as (A) but using affine transformation for the optimal alignment. (D) Same as (A) but for entire skulls and affine transformations.

## 3 Form and feeding mechanics of beaks

Beaks are under multiple selection pressures, and their evolution may be correlated with other parts of the body due to developmental constraints or co-adaptation [15, 16]. Here we explore correlations between beak shape and diet for birds in our study [17]. We grouped species in our dataset by their main diet items following [5]. To explore the correlation between diet and the morphospace generated in the previous sections, we assign each diet category a numerical value 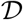, which is an integer in the range [1, 9], and then found the Pearson correlation coefficient between this measure and the morphometric quantity of interest (see Fig. 4). Since the numerical value of 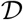 for each diet category involves an arbitrary choice of ordering the categories, we calculate the correlation coefficient for all possible permutations of assigning a value to a diet category and define the correlation coefficient as the maximum across all possible permutations. To check that this method does not generate spurious correlations, we generate control morphospaces consisting of values chosen at random between [0,1], and then compute the correlation between these random values and 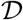. By repeating this for ten thousand trials, we estimate that correlations above 0.25 are statistically significant since it did not result for any of these trials. Using this method, we find that the aspect ratio *W_C_/L_C_* is highly correlated with diet (with coefficient of 0.7), where small values correspond to nectar feeding species and large values correspond to fruit and seed eating species (Fig. 4A), possibly enabling them to exert larger bite forces as described in [18, 19]. Fruit and seed eating species also have high sharpening rate 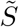 (coefficient ~ 0.5, Fig. 4B), possibly enabling their beak to withstand high forces and increasing fracture resistance [6].

**Figure 4:**
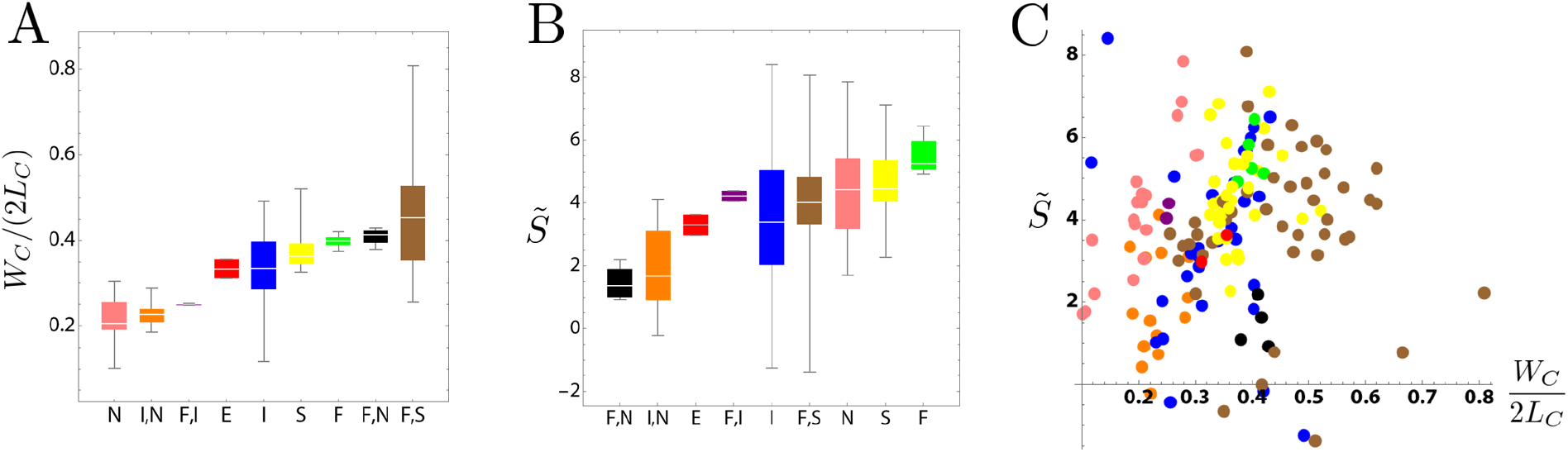
Diet and evolutionary morphospace. (A) For each diet category, we plot the aspect ratio *W_C_*/(2*L_C_*) which is highly correlated with the diet value (each diet category has a value from 1 to 9 in the order shown in the figure). The correlation coefficient is *Corr*[*W_C_*/(2*L_C_*), 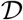] = 0.73. The abbreviations stand for: N = Nectar, S = seeds and nuts, F = fruits and leaves, and E = eggs and blood. (B) Diet category against the sharpening rate 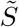, with corresponding correlation coefficient 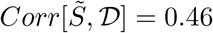. (C) Each specimen is represented as a point in the morphospace (*W_C_*/(2*L_C_*), 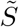), and color coded according to its diet category as given in panel (A) and (B).

## 4 Beak development

Beak growth occurs in group of dividing cells near the tip of the developing beak, and the number of dividing cells in the growth zone diminishes over time [9]. By looking at how adult beak cross sections vary along the centerline, we find that the shape of the cross sections changes, as their size decreases towards the tip (SI Fig. S6). This observation of 3D beak shapes rules out homogeneous and isotropic contraction of the growth zone over time to explain beak shapes [9,14]. To account for the change in cross sections over time, we previously proposed a cellular model that accounts for cell proliferation and movement patterns in space-time, and a coarse-grained tissue-level model for the evolution of the beak treated as a continuum [6]. By modifying this model to account for the centerline curvature of the beaks, we obtain a model that generates all the beak shapes in our data set by varying a single dimensionless parameter (describing how the dynamics of growth depends on curvature), along with the initial size and shape of the cross section (Fig. 5D), the centerline curvature of the beak resulting from continuous turning of cell division plane (orange arrows in Fig. 5D), and the extrusion rate of the growth zone along the proximal-distal axis, denoted as *U*.

**Figure 5:**
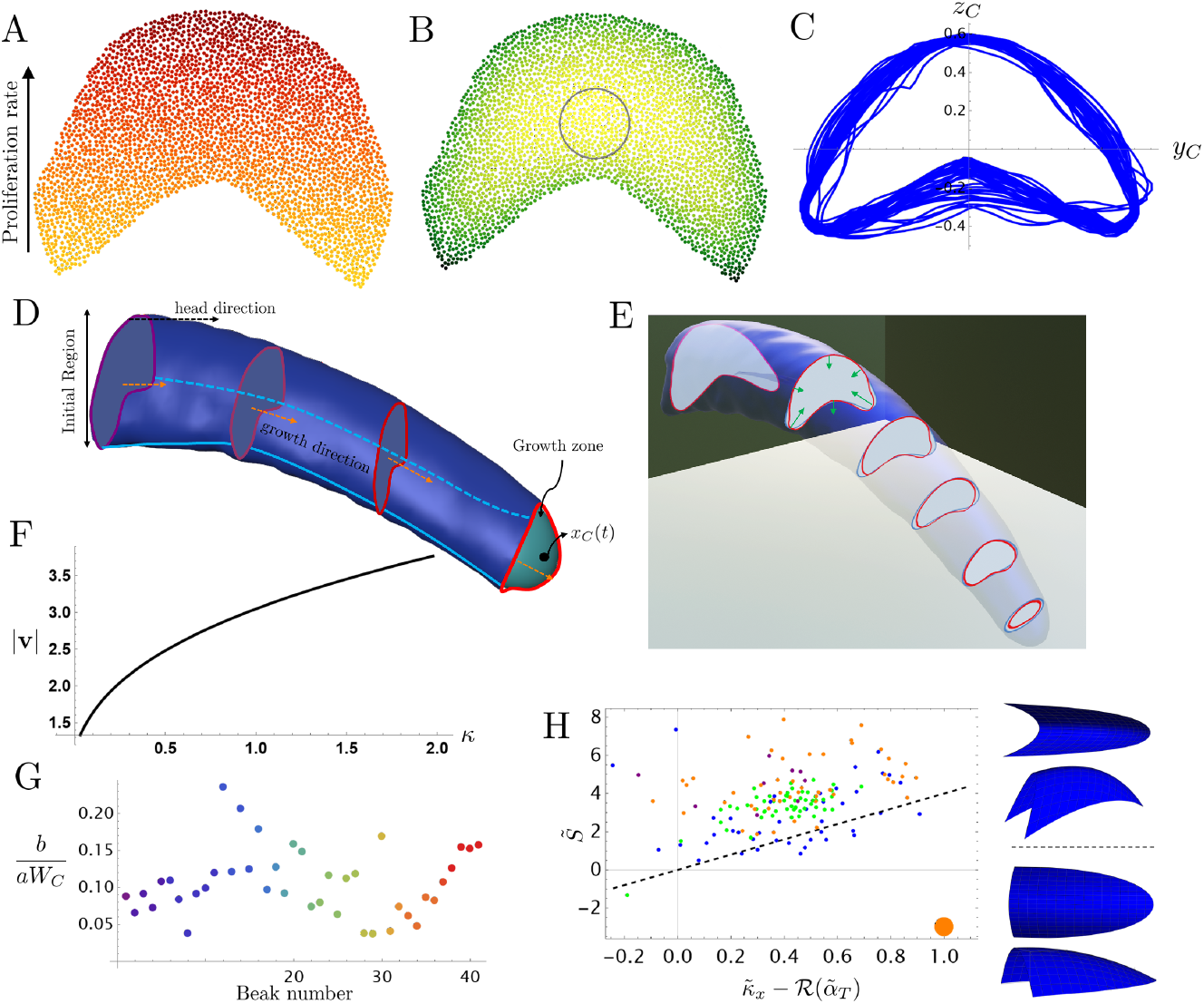
Beak development. (A) The assumed linear gradient in cell proliferation leads to a continuously turning growth direction leading to beak centerline curvature. (B) Morphogen concentrations from the boundary death model described in the main text, the gray circle represents the range of the morphogen. (C) Basal cross-sections from the beaks of all honeycreeper specimens, scaled to have unit width and depth. (D) The blue mesh is taken from the v. *coccinea* sample shown in Fig. 2. The function *x_C_*(*t*) is the center of the growth zone at time *t*, which is the green-blue region near the tip. Also illustrated is the direction of growth (orange arrows) and the major axis of the skull (black arrow). The red curves represent cross-sections at different times, with more recent ones being brighter. (E) Results for the same v. *coccinea* sample as above with cross sections generated from a simulation using the mean curvature flow, **v**= −(*a* + *b κ*)**n**, where **v** is the velocity vector due to the evolving transverse cross sections (green arrows), *κ* is the arc curvature at the corresponding point on the cross section, and **n** is the unit normal to the cross section at that point. The dimensionless parameter used to generate this result corresponds to *b*/(*aW*) = 0.15. The initial conditions were generated from a basal cross section of the actual beak. (F) speed of points on the cross section, taken at *x_C_* = 0.5*L_C_*, as a function of the arc-curvature of the cross section. (G) The dimensionless parameter *b/aW_C_* plotted for all honeycreepers, color coded as in Fig. 1C. (H) The dashed line represents the constraint derived in the main text, 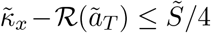, which is approximately satisfied by the honeycreeper samples in our dataset. Here 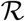(·) ≡ max(0, ·) is the unit ramp function. The two blue meshes (each viewed from two angles) correspond to the orange point, where the upper one has 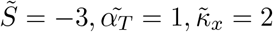 and the lower one has 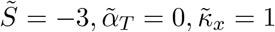.

### 4.1 Cell-scale model for growth

In order to achieve a continuous turning of the growth zone and generate a curved beak, cells must divide and extrude faster towards the upper part of the beak (culmen). To achieve this, we assume a linear gradient in cell proliferation rates (Fig. 5A), achieved for example by gradients in Bmp4, which were shown to help generate the highly curved cockatiel beaks compared to duck and chick in [20]. Furthermore, to account for our observation of changing shapes of cross sections, we employ the model we developed in [6], where cells stop proliferating based on the concentration of another morphogen, produced by cells in the growth zone, that is approximated by a narrow slice of tissue along the proximal-distal axis. When the concentration of the morphogen at some position drops below a threshold, cells stop dividing there. The morphogen diffuses to the surrounding cells with diffusion constant *D_c_* and degrades at a rate Γ. This limits the efficacy of the morphogen produced by a cell to a region of size 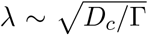 (gray circle in Fig. 5B). Furthermore, since only cells in the growth zone produce this morphogen, a gradient will be established along the proximal-distal axis from which cells can generate a stable growth in the distal direction.

Due the morphogen signaling — whose range is given by 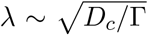 — cells behave differently depending on how far they are from the boundary of the growth zone; cells whose distance from the boundary is smaller than *λ*, will receive morphogen signals from fewer cells. For the same reason, cells proliferate less in more positively curved regions near the boundary (e.g., black parts in Fig. 5B). In addition to this effect, which we term boundary death, there can also be bulk death, where cells stop dividing at some probability *P_death_*, independent of the morphogen concentration. This last effect, when acting alone, leads to an exponential decay over time of the growth zone size, without any change in its shape.

### 4.2 Tissue-scale model for growth

Next, we consider a tissue scale approximation of the cellular model described above. Since cells near a curved part of the growth zone boundary receive the morphogen signal from fewer neighbors (Fig. 5 B), the envelope of the growth zone will shrink faster in those regions. By looking at transverse sections of the adult beaks, represented using Eq. (1), we observe that regions of higher curvature (near *y* = 0) do in fact shrink faster (Fig. 5E). Therefore, this suggests that the boundary of the growth zone evolves with a velocity that depends on the local (arc) curvature through the relation

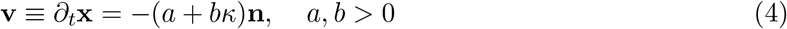

where **n** is the inward normal to the surface. The ratio *b/a* gives a length scale that determines which of the two terms in the growth law will dominate. When the radius of curvature *κ*^−1^ of the transverse beak cross section is greater than *b/a* the term *a***n**, which also describes the evolution of a wavefront under Huygens’ principle, dominates. when *κ*^−1^ ≪ *b/a*, the term *bκ* dominates. This growth law is the well-known curve shortening (or mean curvature) flow in which speed is proportional to the curvature, introduced to account for the erosion of pebbles [21], grain growth from a melt [22].

Fig. 5D shows the agreement between the shape generated by the mean curvature flow and the cross sections of a V. *coccinea* sample (see also SI Fig. S7). By fitting the mean curvature flow to beak cross sections, we obtain the dimensionless parameter *b*/(*aW*), which is plotted in Fig. 5F.

### 4.3 Developmental constraints in morphospace

The existence of a generative model that can reproduce beak shapes and is inspired by the developmental biology of the beak, implies that some beak shapes are not allowed. Beak shapes generated by Eq. (4), when *a* and *b* are both positive, will have cross-sectional area and perimeter that decrease over time (SI Fig. S6A-B) and their cross sections should become more convex and circular as we approach the tip (SI Fig. S6C-D). In this section, we investigate further constraints on beak shape that follow from the curvature driven flow described in Eq. (4).

To find analytic expression for these constraints, we simplify Eq. (4) by assuming circular cross sections with radius *R*(*t*) = *W* (*t*)/2, where *W* (*t*) is the width of the beak as a function of time, which we will calculate using the tomium width given in Eq. (3). The corresponding beak curvature will be *κ*(*t*) = 1/*R*(*t*) = 2/*W* (*t*) and, after plugging this into Eq. (4), we obtain

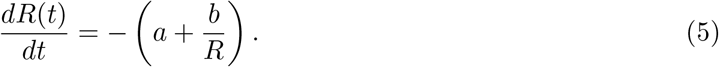

To convert between spatial (*x_C_*) and temporal variables (*t*), we use the extrusion speed *U* so that (*t* − *t*\*)*U* = *x_C_*, where *t*\* = *L_C_/U* is the final time when the cross section shrinks to zero size at the tip of the developed beak. Near the tip (*x_C_* → 0), the term proportional to *b* in Eq. (5) dominates and we get

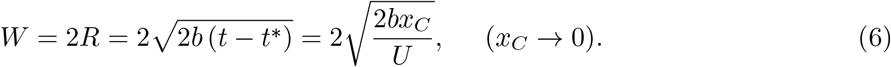

On the other hand, we have found in Sec. 2.1 that the beak width is given by Eq. (3), which leads to the following when (*x* → 0),

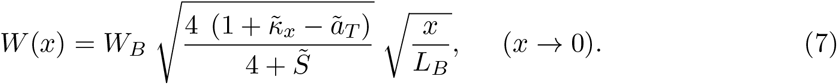

After using the Pythagorean theorem to estimate the factor 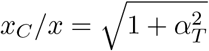 near the tip and equating the right hand sides of Eqs. (6 - 7), we get

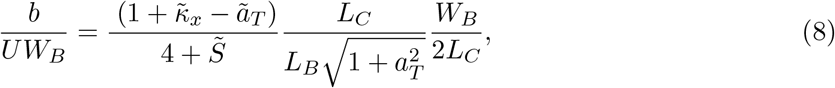

Since the beak shrinks to a point at the tip, we require that *a* > 0, *b* > 0. Therefore, for a given value of *b*, the largest possible length *L_C_* happens when *a* = 0 (since higher value of *a* reduce the length via faster shrinking of the growth zone). When *a* = 0, we have 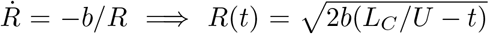. From the initial conditions we get *R*(0) = *W_B_*/2 = 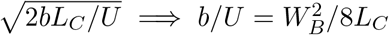. Since larger values of *L* require negative values of *a*, we get the prediction *b*/(*W_B_U*) < *W_B_*/8*L_C_*, which can be rewritten using Eq. (8) to eliminate *b* in terms of other morphological parameters and gives

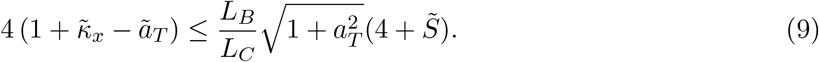

The factor multiplying 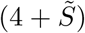 is very close to unity in our dataset and we may simplify the constraint to 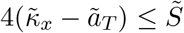, which reduces to the earlier result 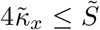, when 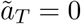. Since all other species satisfy the constraint 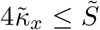 and have *a_T_* ≤ 0 unlike honeycreepers (Fig. 2 and SI Fig. S3B), we introduce the ramp function 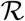(·) ≡ max(0, ·) and write the constraint as 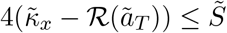. Figure 5G shows that honeycreepers move closer to the region predicted by this constraint, which may be compared with two examples of beak shapes that would not be possible due to this constraint (blue meshes in Fig. 5G).

## 5 Conclusions

Rather than using landmark data as usually done, this paper develops novel morphometric approaches to elucidate the three-dimensional beak morphology and link evolution, mechanical function, and development. Building on our prior study of Darwin’s finches [6], we found geometric functional forms, Eq. (1), that capture the beak shape of samples in our dataset, including Hawaiian Honeycreepers (HC), Darwin’s finches (DF), and their respective relatives by varying a small number of geometrically meaningful parameters — orientation relative to the skull, aspect ratios and curvatures (Fig. 2). The highly curved HC beaks (Fig. 1) necessitated developing a method to extract the curved beak centerline and its perpendicular cross-sections (Fig. 2C and SI Fig. S2D-E).

Our analysis uncovered interesting patterns in morphospace, including that HC beaks occupy a broader region of morphospace, a finding we validated through employing parameter-free, mesh-based, methods using the Hausdorff distance and multi-dimensional scaling (Fig. 3). The mesh-based comparative analysis also showed that DF beaks occupy a distinct region of morphospace (Fig. 3A), even with affine transformations that lead to overlap in regions occupied by other taxa (Fig. 3C). Surprisingly, we found that some highly curved beaks, such as those of *V. coccinea*, do not have a high value of centerline curvature in absolute units, but rather have a high value of the (scale invariant) quantity curvature times length 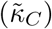. We also explored the relation between beak shape and diet and find a correlation that we can rationalize by considering the beak as a mechanical tool for feeding [19, 23, 24]. In particular, the width to length aspect ratio and the sharpening rate (how fast the curvature of transverse sections increases towards the tip) are highly correlated with diet.

To further understand the source of variation in HC beak morphology, we employed developmental models that use adult beak shape to glean information about beak ontogeny. While assuming a single growth direction, parallel to the major axis of the beak, was sufficient to generate DF beaks [6], generating HC beaks required a changing growth direction by assuming a linear gradient in cell proliferation rates (Fig. 5). With this modifications, our developmental model explains how beak shape emerges due to initial size, shape, growth direction relative to the skull, the extrusion velocity of the growth zone, and the transverse shrinkage rate determined by both *a* and *b* parameters. Remarkably, a single dimensionless parameter, *b*/(*aW*), determined how cross sectional shape evolves along the centerline for all species in our dataset.

The fact that honeycreepers occupy a broader range of morphospace in our analysis, and required a modification of our developmental model to account for centerline curvature, raises the question of how this group managed to escape the constraint that limited the variation of other birds in our dataset. Applying the methods developed in this paper in conjunction with experimental work on developing beaks to a wider range of avian taxa would be natural next steps towards a more comprehensive description of beak shaping during development, its evolution across time, and its function as a remarkably adaptable tool.

