## Supplementary material for "Beak morphometry and morphogenesis across avian radiations": SI

#### S1 Dataset

Our dataset contains 151 specimens that were collected and scanned using X-ray microcomputed tomography, as described in [1], to obtain the 3D triangulated mesh of the full skull (Fig. S1). The scans — which are divided into Hawaiian Honeycreepers (n = 41), their relatives (n = 9), Darwin's Finches (n = 54), and their relatives (n = 47) — are handled using the open-source software Blender [2], which was used to cut beak from the skulls.

#### S2 Mesh processing and feature extraction

To separate the beak from the skull and analyze their morphologies separately, we rotate the skull to align the forward direction with the $x-$axis and use a cutting plane that captures as much of the upper surface of the beak as possible (shown in orange in Fig. S2A).

Once we have both the raw full skull scans and the raw beaks, we convert each of them into a smooth and watertight (genus-0) triangulated mesh using the mesh processing method described in [3,4], which involves a voxelization step using the MATLAB custom function `Polygon2Voxel` [5] and the built-in functions `alphaShape` and `boundaryFacets`, a hole filling step using a medial axis approach [6], a smoothing step using the implicit Laplacian smoothing [7] with the cotangent Laplacian [8], and a repairing step using the Iso2mesh toolbox [9] (see Fig. S2B-C). To improve the approximation accuracy of the watertight mesh, we further align it with the raw mesh using the Iterative Closest Point (ICP) method [10] and apply an anisotropic rescaling based on the size of the two meshes along the $x, y, z$-directions.

Next, we develop an iterative method for extracting optimal centerlines and cross sections from every watertight beak mesh. We start by finding the intersections between the mesh and equally spaced vertical planes (with $x = constant$) using the MATLAB custom function `intersectPlaneSurf` [11], which gives a set of vertical cross sections in the form of closed curves. By computing the centroids of all cross section curves and connecting them, we obtain an initial centerline (see Fig. S2D, left). Because of the curved geometry of the beaks, the vertical cross sections and the centerline may not be an optimal representation of the beak shape. Now, we compute a new set of intersections between the mesh and planes normal to the centerline at equally spaced points. As the normal direction of the centerline changes with the curvature of the centerline, the new cross sections represent the beak shape more accurately. We can then use the centroids of the new cross sections to form an updated centerline curve (see Fig. S2D, middle). By repeating the above procedure, we obtain an optimal centerline and an optimal set of cross sections for our analysis (see Fig. S2D, right). Fig. S2E shows 2D views of sampled cross-sections from a *V. coccinea* sample.

### S3 Parametric morphometrics

In this section we describe how we extract the parameters of that describe the beak shape. By parametric morphometrics, we mean that the shape is fit using a parameterized family of functional forms and the resulting parameters give geometrically interpretable measure (such as curvature) of beak shape.

As described in the main text, we extract the upper beak by finding the points having the maximum and minimum $y$-values, which we assume correspond to the tomium edge shown in Fig. S3. We then align the resulting points to their moment directions by computing the largest eigendirection of the covariance matrix (major axis) and rotating the mesh around the y-axis (which perpendicular to the plane of symmetry of the beak) so that the major axis lies as close as possible to the x-axis. Once the axes are aligned we fit the resulting points to the form given in Eq. (2.1) of the main text. Since some meshes have pieces missing near the tip, we allow the tip position to be inferred using the best value of the following form

$$z_U(x,y) = z_{tip} + a_U\ (x - x_{tip}) - \kappa_x\ (x - x_{tip})^2 - (\kappa_{tip} - S\ (x - x_{tip}))\ (y - y_{tip})^2. \tag{S.1}$$

The dimensions of the beak, denoted as $(L_B, W_B, D_B)$ are found by finding the minimum bounding box (aligned to the moment directions). Note that the value of nondimensionalized parameters, will depend on how much of the beak is included in the cutting procedure and in normal (to the centerline) cross sections obtained as described in the previous section. When comparing between beaks it is important to standardized this procedure.

### S4 Surface-based morphometrics

As described in the main text, we can perform a multidimensional scaling (MDS) analysis by optimally aligning every pair of beak or skull surface meshes. More specifically, we search for the optimal transformation among a prescribed class of transformations to minimize the symmetric Hausdorff distance between the pair of surface meshes. Different classes of transformations can be considered:

1. Rigid transformations (i.e. a combination of translation and rotation).
2. A combination of translation, rotation and isotropic scaling.
3. A combination of translation, rotation and anisotropic scaling.
4. Affine transformations (i.e. a combination of translation, rotation, scaling, and shear).

In the main text, we presented the MDS analysis based on rigid and affine transformations. In Fig. S8, we also show the MDS results for the beaks using the two other classes of transformations. It can be observed that by introducing isotropic rescaling (Fig. S8A), the range of variation in the morphospace for each group is reduced. Also, the region of morphospace occupied by Darwin's finches remains distinct, while the overlap of the regions for the other three groups becomes more significant, which suggests that part of their shape variation can be explained by a uniform rescaling. By further allowing the rescaling factors for the three axes to be different in the alignment transformations (Fig. S8B), we can see that the overlap for the three latter groups is further increased, which suggests another part of their shape variation is related to their relative size in different dimensions. Finally, introducing the use of affine transformations yields a similar result (Fig. S8C), which suggests that shearing is not a significant factor in the variation of the beak shapes.

Fig. S9 shows the additional MDS results for the full skull shapes. It can be observed that even after introducing isotropic scaling (Fig. S9A), anisotropic scaling (Fig. S9B) and affine transformations (Fig. S9C), the overlap between the regions occupied by the different groups remain significant.

#### S5 Fitting the parameters of the modified mean curvature flow

We simulate the modified mean curvature flow in two ways. The first uses the level set method as described in [4,12] and was used to generate the results in Fig. S7. Since this method involves long computation time and fitting parameters was by trial and error, we employed a faster method to automatically fit the results of all beaks. We numerically compute the curvature and normal (after employing a low-pass filter to smooth the data) using a cubic spline with periodic boundary conditions, which is provided in the package [13]. Then using this information the cross section is updated according to the equations of motion (Eq. (4.2) in the main text). To avoid numerical instabilities as the cross sections shrinks to smaller size, the cross section is resampled (using the cubic spline) to keep the density of points constant and uniform.

By rescaling time to $\tilde{t} = t/a$ and lengths $\tilde{\mathbf{x}} = \mathbf{x}/W_C$, we obtain

$$\tilde{\mathbf{v}} \equiv \partial_t \tilde{\mathbf{x}} = -\left(1 + \frac{b}{aW_c}\kappa\right)\mathbf{n}. \tag{S.2}$$

Therefore, the evolution of the cross-sections is controlled by a single dimensionless parameter. To find the optimal value of this parameter for each beak, we quantify the fit by comparing each simulated cross section to the corresponding cross section from the beak using a Hausdorff distance. The correspondence between simulation time and beak cross section is done relative to the total length of the centerline, or total time of the simulation, so that $\tilde{t}/\tilde{t}_{final} = x_C/L_C$.

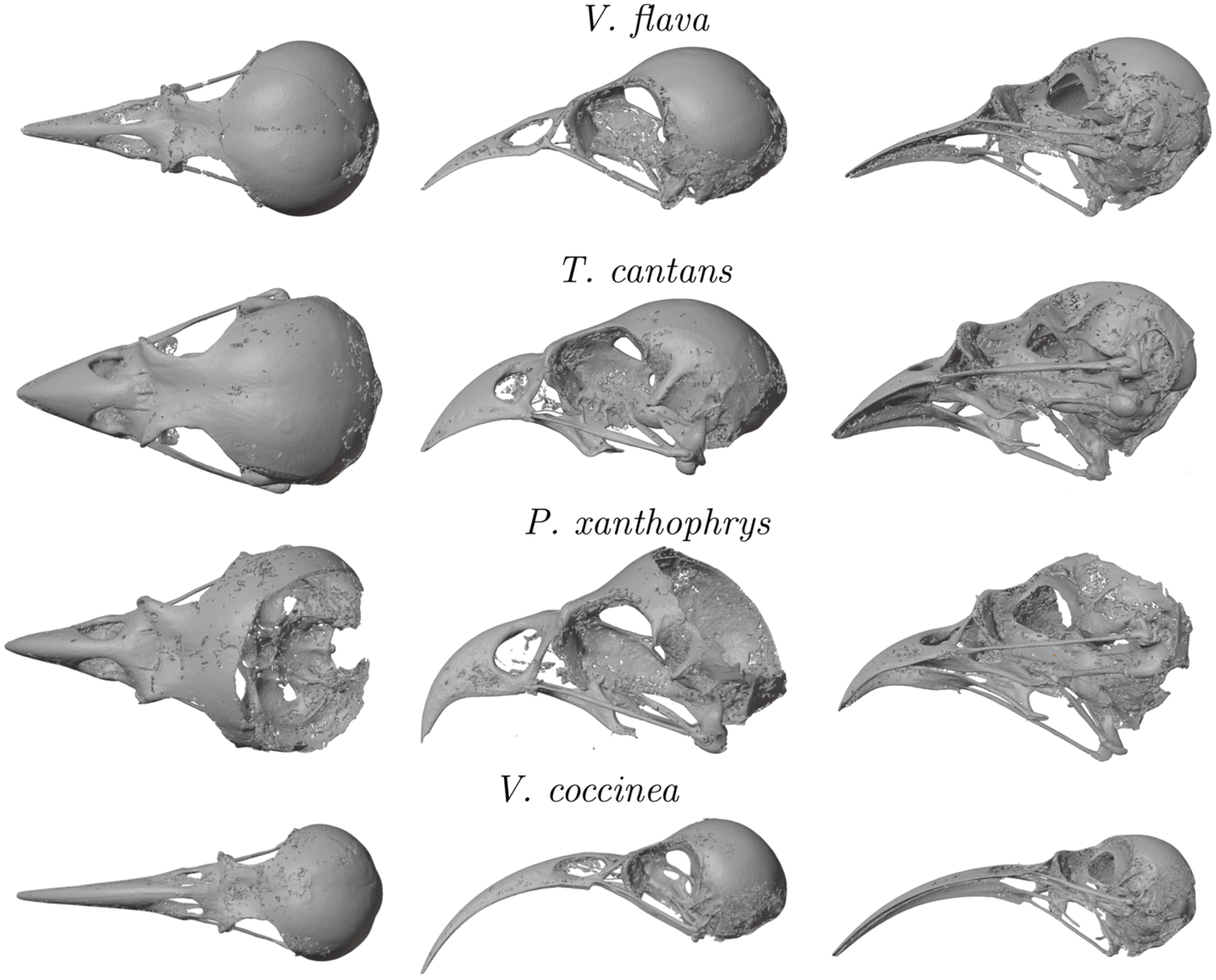


Figure S1: **Dorsal, lateral, and ventral-lateral views of some of the honeycreeper skulls used in our study.**

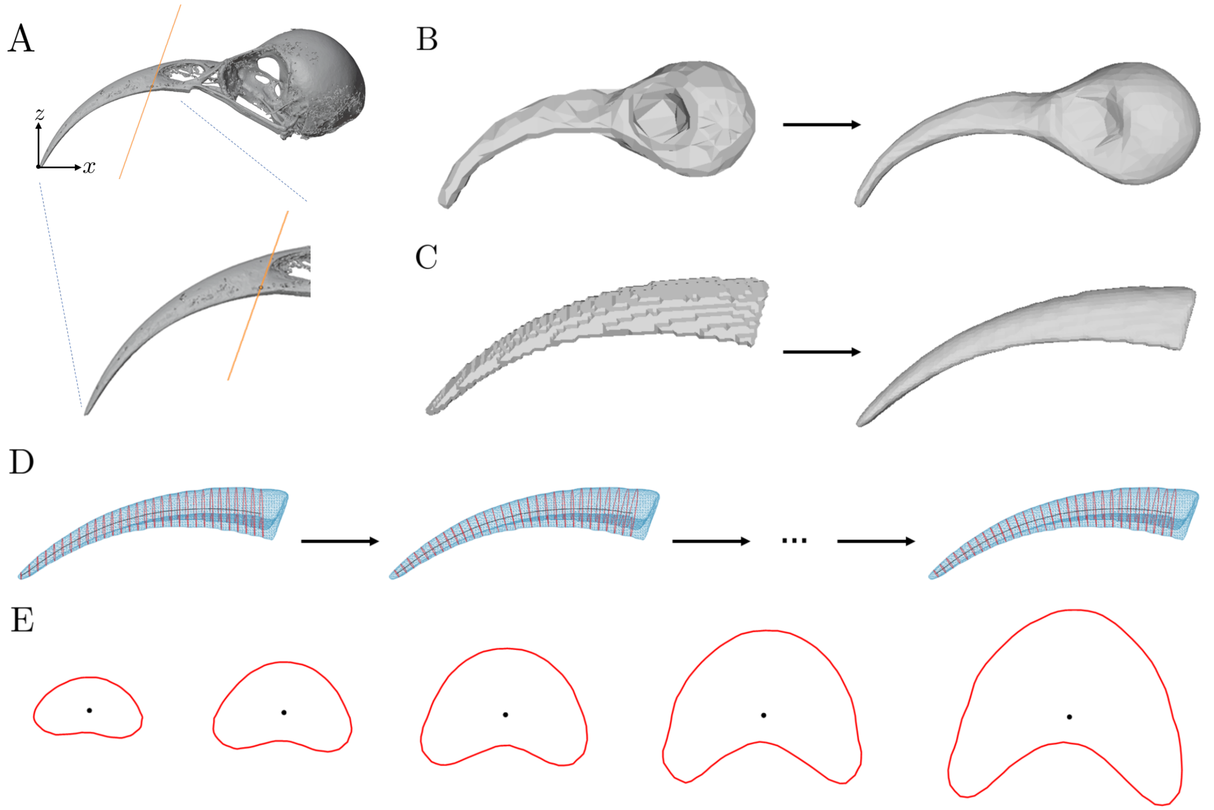


Figure S2: **Processing the 3D scans and extracting the beak centerline.** (A) A projected view of the skull onto the $xz$-plane. The skull is aligned (in Blender [2]) so that the x-axis is aligned as shown and the orange plane represent the cut performed to separate the beak from the skull. (B) The voxelization and smoothing of a full skull surface. We first apply a voxelization step to remove the holes in the raw 3D scan, and then apply a smoothing step to improve the result. (C) The voxelization and smoothing a beak surface. (D) Extracting the cross-sections and centerlines of a beak surface via an iterative algorithm. In each step, we compute the cross sections and then extract their centroids to form a centerline. We then use a new set of planes normal to the centerline to compute a new set of cross sections, and repeat the above process to eventually get an optimal set of cross sections and an optimal centerline. (E) The extracted 5th, 10th, 15th, 20th, 25th cross-sections in the result in (D).

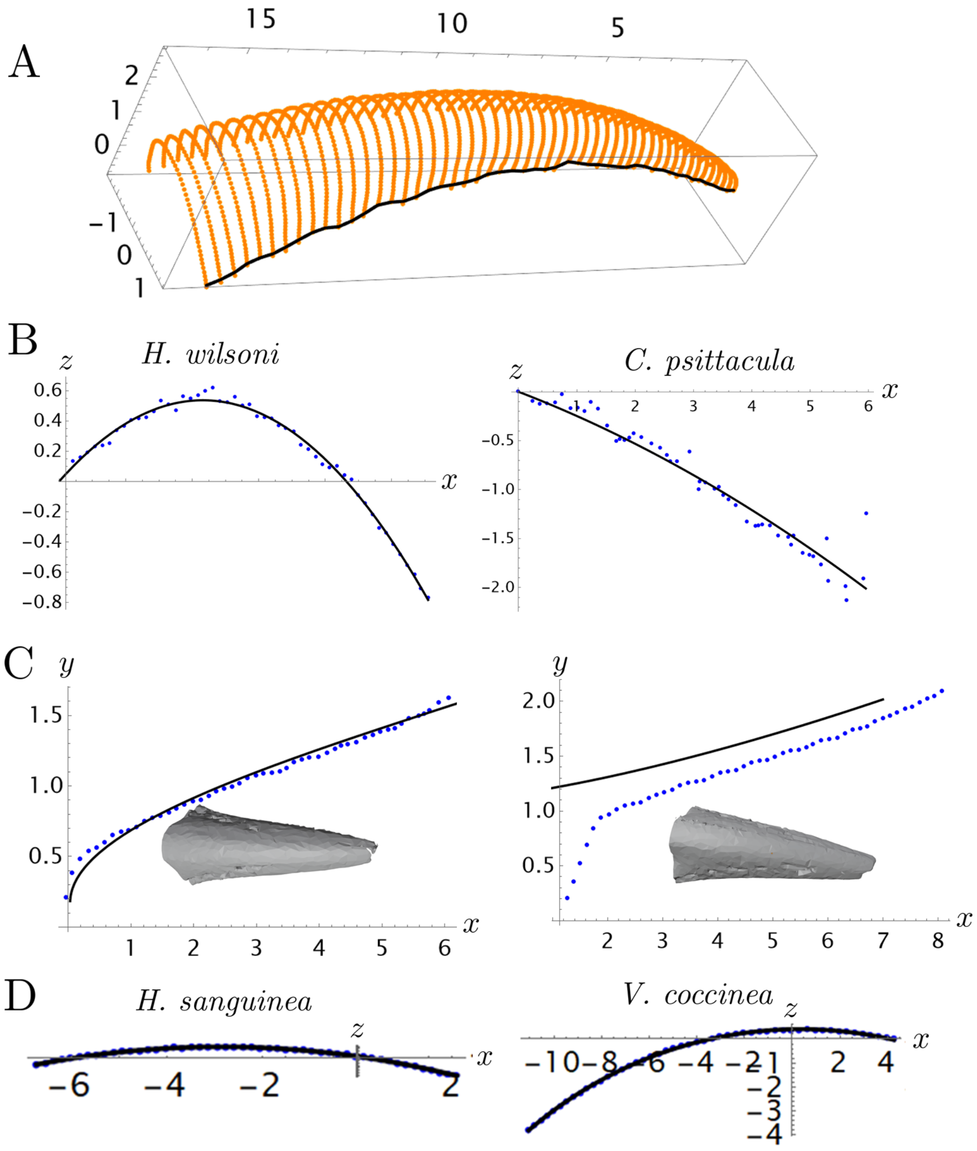


Figure S3: **Processing the beak cross sections.** (A) The extracted upper beak and tomium points. (B) Parabolic fit ($z_T = a_T x - \kappa_T x^2$) of the tomium points, when projected onto the $xz$-plane. (C) The beak width given by Eq. (2.3) in the main text and the projection of the extracted beak tomium onto the $xy$-plane. (D) Extracted beak centerlines and corresponding circular fits, which are in very close agreement for almost all specimens.

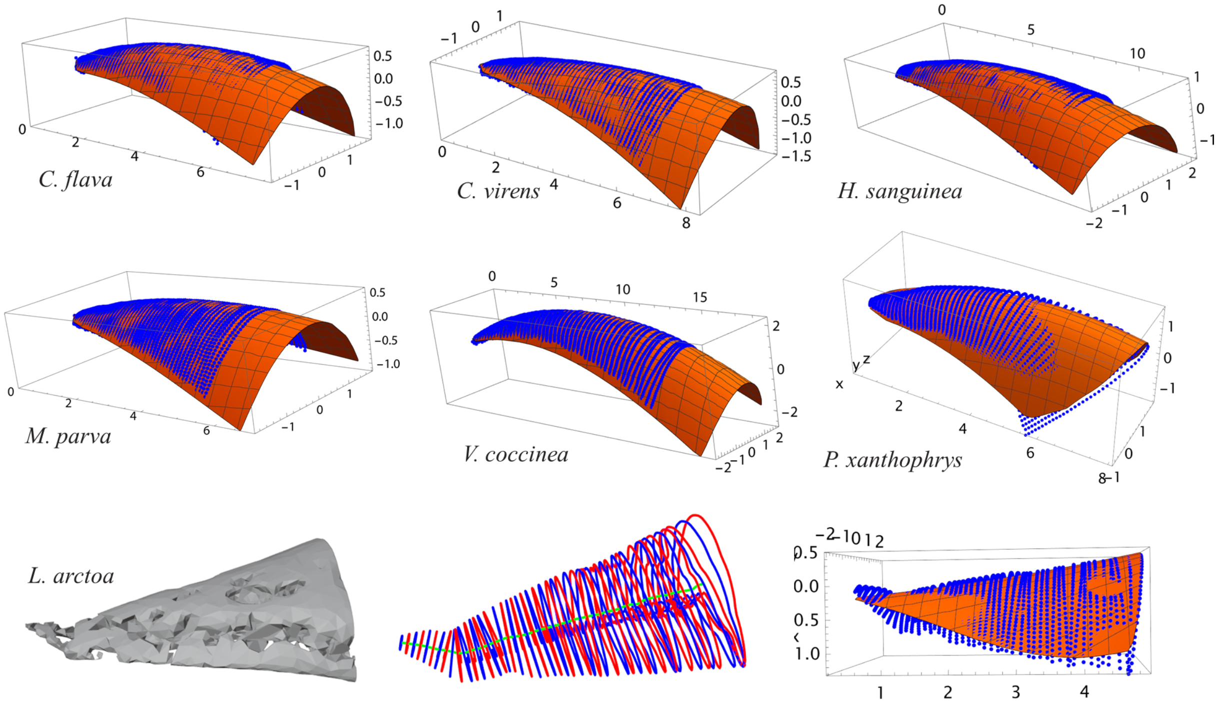


Figure S4: **Beak fits using the parabolic form (Eq. (2.1) in the main text).** The fits are in reasonable agreement with the blue points that correspond to the upper part of the beak mesh where the coordinate axes are aligned with the beak moment directions. Due to imperfections in the scan, *L. arctoa* exhibits a negative sagittal curvature.

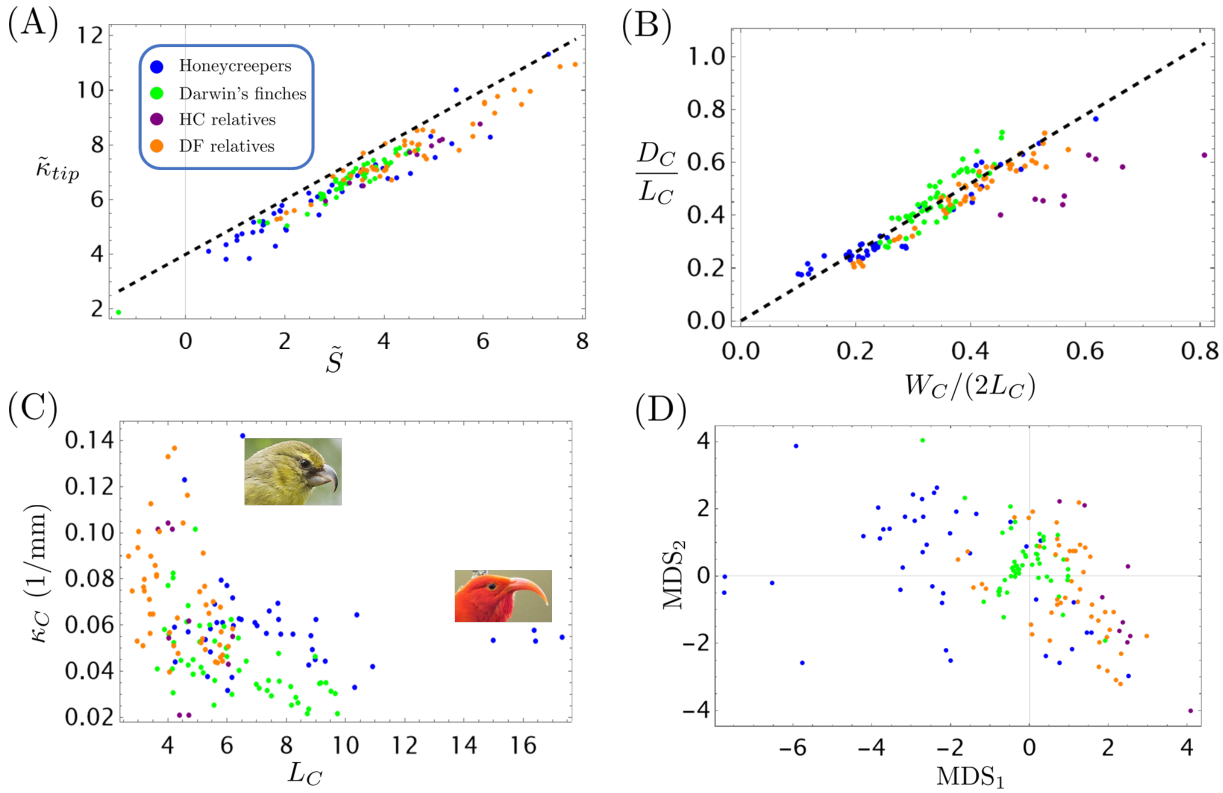


Figure S5: **Beak morphospaces.**

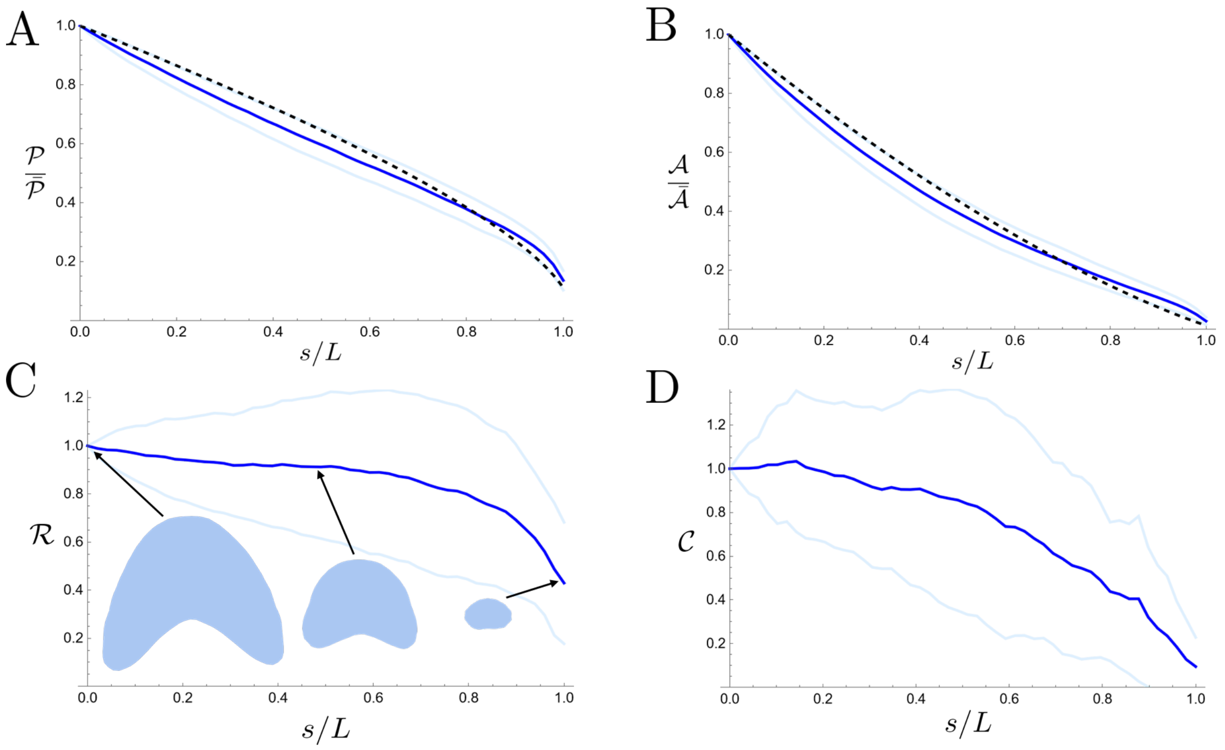


Figure S6: **Cross sections change shape along the length of the beak.** (A) Plot of transverse cross-sectional perimeter $\mathcal{P}$ as a function of normalized arc-length $s/L$, where $L$ is the length of the centerline. The blue curve gives the average over the Honeycreeper specimens, with the standard deviation indicated by the confidence bands. The dashed black curve gives the mean curvature flow fit. (B) Corresponding plot of area $\mathcal{A}$ versus normalized arc-length. (C) Plot of roundness, defined as $\mathcal{R} \equiv 1 - 4\pi\mathcal{A}/\mathcal{P}^2$, (D) Plot of convexity, defined as $\mathcal{C} = 1 - \mathcal{P}_{\mathcal{C}}/\mathcal{P}$, where $\mathcal{P}_{\mathcal{C}}$ is the perimeter of the convex hull of the cross section.

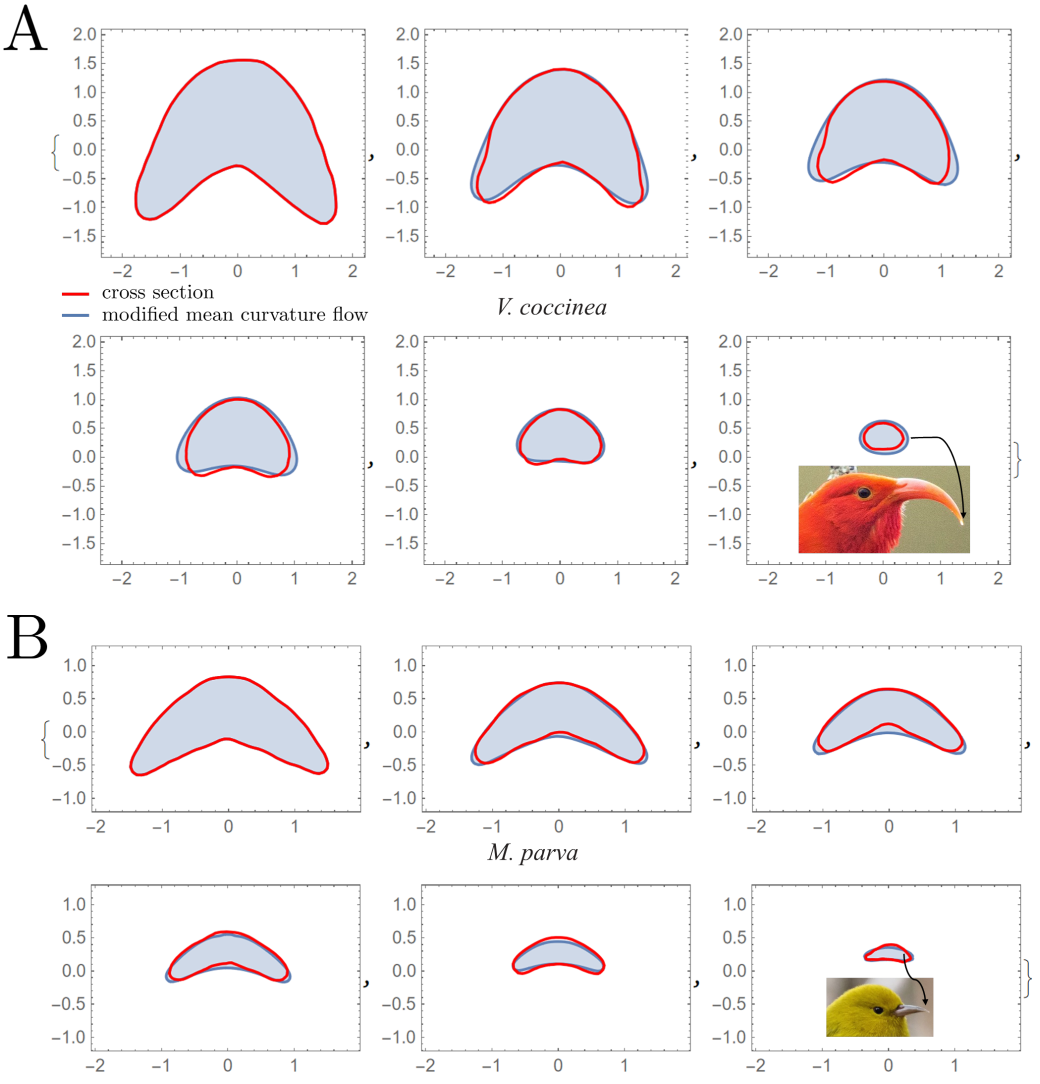


Figure S7: **Simulation results using the modified mean curvature flow, compared with extracted normal beak cross sections for (A) *V. coccinea* and (B) *M. parva*.**

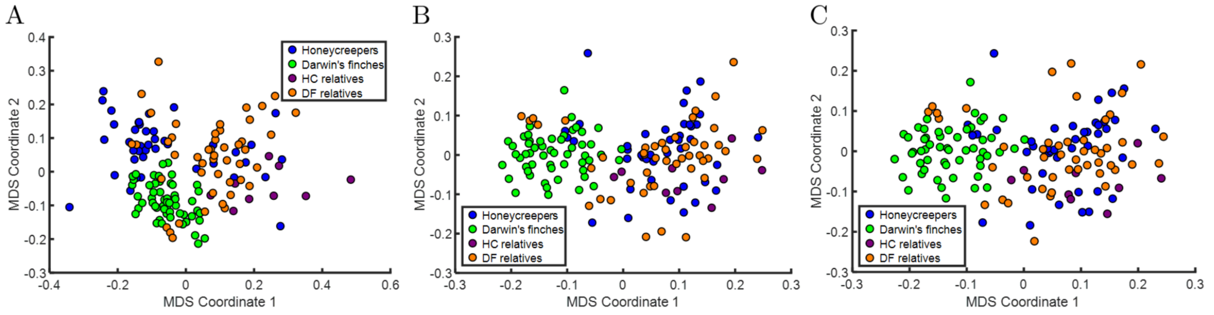


Figure S8: **Multidimensional scaling (MDS) analysis for the beak shapes.** (A) The result obtained by optimally aligning the beak shapes using a combination of translation, rotation and isotropic scaling. (B) The result obtained by optimally aligning the beak shapes using a combination of translation, rotation and anisotropic scaling. (B) The result obtained by optimally aligning the beak shapes using affine transformations (a combination of translation, rotation, scaling and shear).

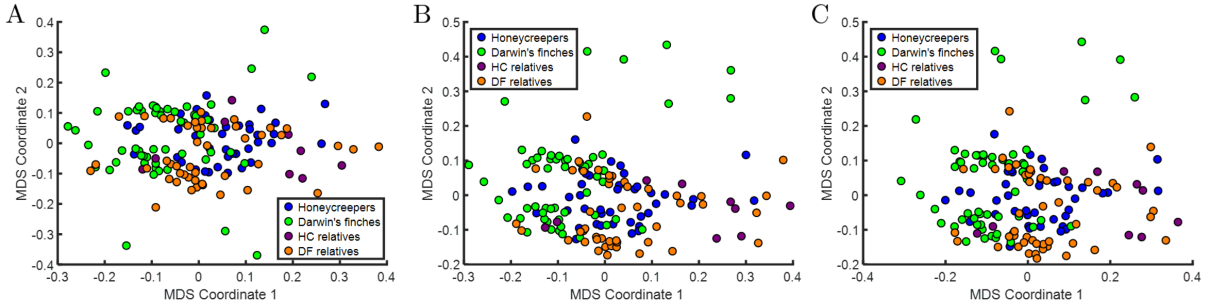


Figure S9: **Multidimensional scaling (MDS) analysis for the skull shapes.** (A) The result obtained by optimally aligning the skull shapes using a combination of translation, rotation and isotropic scaling. (B) The result obtained by optimally aligning the skull shapes using a combination of translation, rotation and anisotropic scaling. (B) The result obtained by optimally aligning the skull shapes using affine transformations (a combination of translation, rotation, scaling and shear).
